# Influenza A virus undergoes compartmentalized replication *in vivo* dominated by stochastic bottlenecks

**DOI:** 10.1101/2021.09.28.462198

**Authors:** Katherine A. Amato, Luis A. Haddock, Katarina M. Braun, Victoria Meliopoulos, Brandi Livingston, Rebekah Honce, Grace A. Schaack, Emma Boehm, Christina A. Higgins, Gabrielle L. Barry, Katia Koelle, Stacey Schultz-Cherry, Thomas C. Friedrich, Andrew Mehle

## Abstract

Transmission of influenza A viruses (IAV) between hosts is subject to numerous physical and biological barriers that impose genetic bottlenecks, constraining viral diversity and adaptation. The presence of bottlenecks within individual hosts and their potential impacts on evolutionary pathways taken during infection and subsequent transmission are poorly understood. To address this knowledge gap, we created highly diverse IAV libraries bearing molecular barcodes on two independent gene segments, enabling high-resolution tracking and quantification of unique virus lineages within hosts. Here we show that IAV infection in lungs is characterized by multiple within-host bottlenecks that result in “islands” of infection in lung lobes, each with genetically distinct populations. We performed site-specific inoculation of barcoded IAV in the upper respiratory tract of ferrets and tracked viral diversity as infection spread to the trachea and lungs. We observed compartmentalized replication of discrete barcoded populations within the lobes of the lung. Bottlenecks stochastically sampled individual viruses from the upper respiratory tract or the trachea that became the dominant genotype in a particular lobe. These populations are shaped strongly by founder effects, with no evidence for positive selection. The segregated sites of replication highlight the jackpot-style events that contribute to within-host influenza virus evolution and may account for low rates of intrahost adaptation.

## INTRODUCTION

The constant evolution of influenza viruses results in recurring seasonal epidemics and has the potential to initiate pandemics in the human population. Viral evolution occurs on multiple scales. Influenza A virus evolves rapidly on the global scale where population-level immunity positively selects new antigenic variants, necessitating reformulation of the seasonal influenza vaccine^1–3^. Yet, on smaller scales, variants with a predicted fitness advantage rarely rise to high frequency within a host or transmit to a recipient, even in the face of vaccine-induced immunity^4–7^. Within-host variation is low and genetic drift plays a large role^8^. To reconcile these observations, it is currently thought that positive selection drives more deterministic processes at the global scale while genetic drift shapes stochastic processes during local transmission^9^.

Genetic drift of influenza virus between hosts is dominated by bottlenecks, neutral processes in which a small number of viral particles found a new infection, often resulting in dramatic reductions in the effective population size. In natural human infections, bottlenecks during airborne transmission of influenza A virus are exceedingly tight, with as few as 1-2 genomes founding infections in the recipient^8^. Similar results have been detected in animal models, where a very small proportion of intrahost variants successfully found new infections^10^. Under these conditions, genetic drift is expected to have a large impact, where diversity in the transmitted population is dramatically decreased with potential loss of beneficial variants as the population transits the narrow bottleneck^11^. Repeated experimental bottlenecks severely restrict viral fitness, and it has been suggested that transmission bottlenecks restrain jumps across host species and the global rate of influenza virus evolution^8,12,13^.

Influenza virus infection occurs in heterogeneous cell populations within complex anatomical structures in a host^14^. Anatomical structures may help establish local sites with high multiplicity of infection that impact reassortment and complementation. Initial sites of infection are influenced by the tissue-specific distribution of sialic acid receptors and their topology^15–17^. Influenza virus replication can occur throughout the respiratory tract, but recent evidence suggests discrete locations contribute virus that is transmitted. Viruses in the upper respiratory tract, specifically those replicating in the soft palate or nasal epithelial cells, contribute most to the population that is transmitted in animal models^10,18,19^. Thus, it is the diversity at the site of transmission that is likely the most important for onward evolution. How viral movement in the respiratory tract and potential compartmentalization affect within-host evolution and ultimately transmission is poorly understood.

A clearly defined intrahost population structure is required to accurately model and predict influenza virus evolution and onward transmission of variants. Although error-prone genome replication generates influenza A viruses with distinct genotypes, the relatively low levels of naturally occurring variation within hosts do not provide sufficient information for fine-grained analysis of population dynamics. We overcame this limitation by introducing a neutral barcode of 10 random nucleotides into two segments of the influenza virus genome and creating rich viral populations with ∼0.6-3×10^5^ uniquely quantifiable members. Using these barcoded viral populations, we capture soft selective sweeps in cell culture and show adaptive changes arose independently multiple times, yet only one lineage became dominant. Infection in ferrets revealed highly compartmentalized replication as virus migrated from the upper respiratory tract to the lung. Bottlenecks between sites led to stochastic sampling of individual viruses from the upper respiratory tract or the trachea that became the dominant lineage in lung lobes, while there was no evidence of positive selection. Thus, viruses infecting the lung do not constitute a large homogeneously mixed population, but rather multiple isolated populations that each undergo bottlenecking events that can severely constrain population diversity and the potential for selection of fit variants.

## RESULTS

### Large scale incorporation of barcodes avoids artificial bottlenecking of variants

Naturally occurring genetic diversity in influenza virus populations is poorly suited for high-resolution characterization of population-level dynamics and cannot accurately enumerate the full spectrum of individuals present. Moreover, genetic mutations may affect the fitness of the virus, biasing representation of specific variants in the population and precluding their use as a neutral marker. To better resolve and quantify the dynamics of the IAV population, we introduced dual molecular barcodes into the genome of the influenza virus isolate A/California/07/2009 (H1N1; CA07) to create individual viruses that are uniquely recognizable and quantifiable via deep sequencing **(Fig 1A)**. Barcodes of 10 randomized nucleotides were introduced onto the *HA* and *PA* segments that were subsequently used for high-efficiency virus rescue **(Fig 1B)**. HA is under selective pressure as the viral attachment protein and principal target of neutralizing antibodies. We might therefore expect barcodes embedded in *HA* to increase or decrease in frequency as a result of selection on this segment. Conversely, PA is subject to less intense selection pressure, and barcodes on this segment may be expected to better represent total population size. For both *HA* and *PA*, barcodes were encoded between the ORF and the UTR. Packaging and bundling signals were duplicated downstream of the barcode to ensure proper gene replication and virion formation^20,21^. The utility of the system was further increased by using the PA reporter construct that co-transcriptionally expresses Nanoluciferase (PASTN)^20^. Finally, the *HA* segment encodes an additional “registration mark,” six nucleotides creating either NheI or PstI restriction sites, that allows us to index and identify separate libraries of *HA* variants.

**Figure 1.**
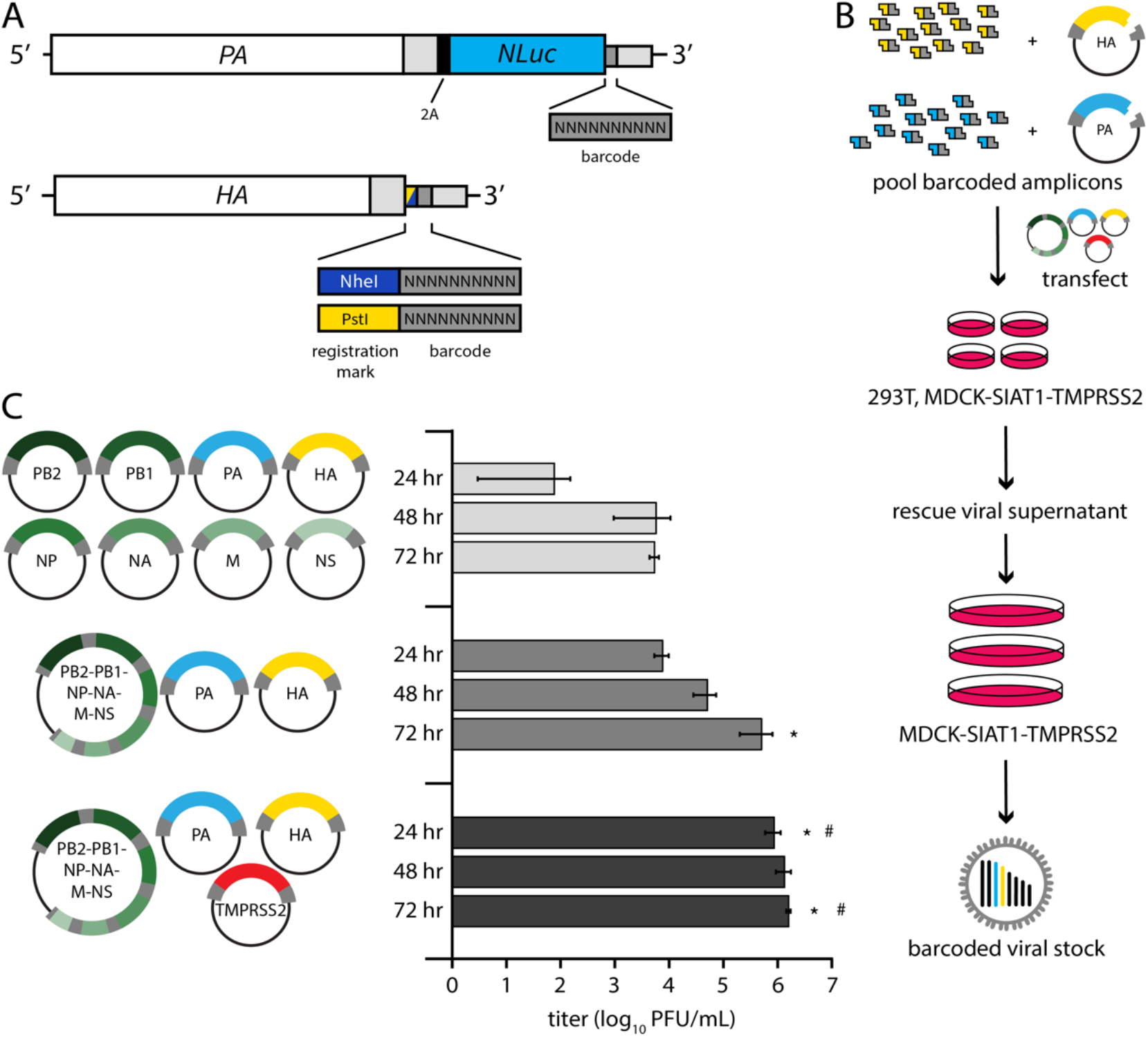
Creation of molecularly barcoded influenza A virus populations. A) Molecular barcodes containing 10 randomized nucleotides were encoded downstream of the open reading frame in the *PA* and *HA* genes, shown as cRNAs. A registration mark was also added onto *HA* to distinguish unique barcode libraries. Sequences were repeated downstream of the barcode to maintain contiguous packaging signals required for replication, and silent mutations were introduced into the open reading frame to avoid direct repeats. B) Experimental overview where randomized barcodes were cloned into reverse genetics vectors followed by large-scale, parallel virus rescues to ensure unbiased barcode distribution. C) Optimized rescue plasmids enhance viral yield. Rescue efficiency was determined by measuring viral titers at the indicated times post-transfection with the standard 8-plasmid system, a consolidated 3-plasmid system, or the 3-plasmid system plus a vector expressing TMPRSS2. (data presented as mean of n = 3 ± sd. ANOVA with Tukey’s post-hoc, * = p<0.05 relative to 8-plasmid rescue, # = p<0.05 relative to 3-plasmid system.)

The limited efficiency of virus rescue can introduce artificial bottlenecks^22^. To increase efficiency and recover a larger, more diverse barcode population, we reduced the number of individual plasmids needed for virus rescue^23^. We combined the 6 non-barcoded segments of CA07 IAV onto a single plasmid, reducing the entire reverse genetics system from 8 to 3 plasmids **(Fig 1C)**. Rescue titers of the 3-plasmid system increased > 200-fold compared to the 8-plasmid system. Titers were further increased by co-expressing transmembrane protease serine 2 (TMPRSS2) during virus rescue, which cleaves and activates HA. We performed 120 parallel virus rescues, pooled the resultant supernatants, and passaged them at high representation to ensure unbiased barcode distribution (**Fig 1B**). NheI and PstI registration marked libraries were prepared independently.

### Viral barcodes reveal selective sweeps *in vitro*

Deep sequencing revealed ∼180,000 unique *PA* barcodes and over 238,000 *HA* barcodes in each of our original virus libraries **(Fig 2A)**. Given that each library has an invariant registration mark, we could measure the fidelity of the quantification pipeline by assessing the number of reads that do not match the predicted registration mark sequence. Over 99% of mapped reads perfectly matched the appropriate registration marks on *HA*, with the majority of those that did not match differing by a single nucleotide from the intended registration mark, indicating the neutrality of the registration mark sequence and overall high fidelity of our sequencing pipeline. Viral populations were characterized by richness, the total number of unique lineages present, Shannon’s diversity, a measure that considers both the presence and relative abundance of a lineage, and evenness, a parameter that compares the frequencies of all lineages in the population to that of a theoretically evenly distributed population^24–26^. Barcode enumeration for viruses recovered from the transfected cell supernatant (i.e. passage 0, P0) revealed rich and highly diverse populations with evenly distributed lineages (**Fig 2A-B**). However, a single passage to create amplified P1 stocks resulted in highly skewed populations in which individual lineages dominated the population, evidenced by dramatic drops in Shannon’s diversity and evenness (**Fig 2B**). The dominant barcodes in each library represented 64-77% of the population. Neither of these barcodes was dramatically over-represented in the plasmid or P0 stocks. In fact, the most abundant lineages in the P0 rescue stocks did not become the most prevalent in our amplified stocks.

**Figure 2.**
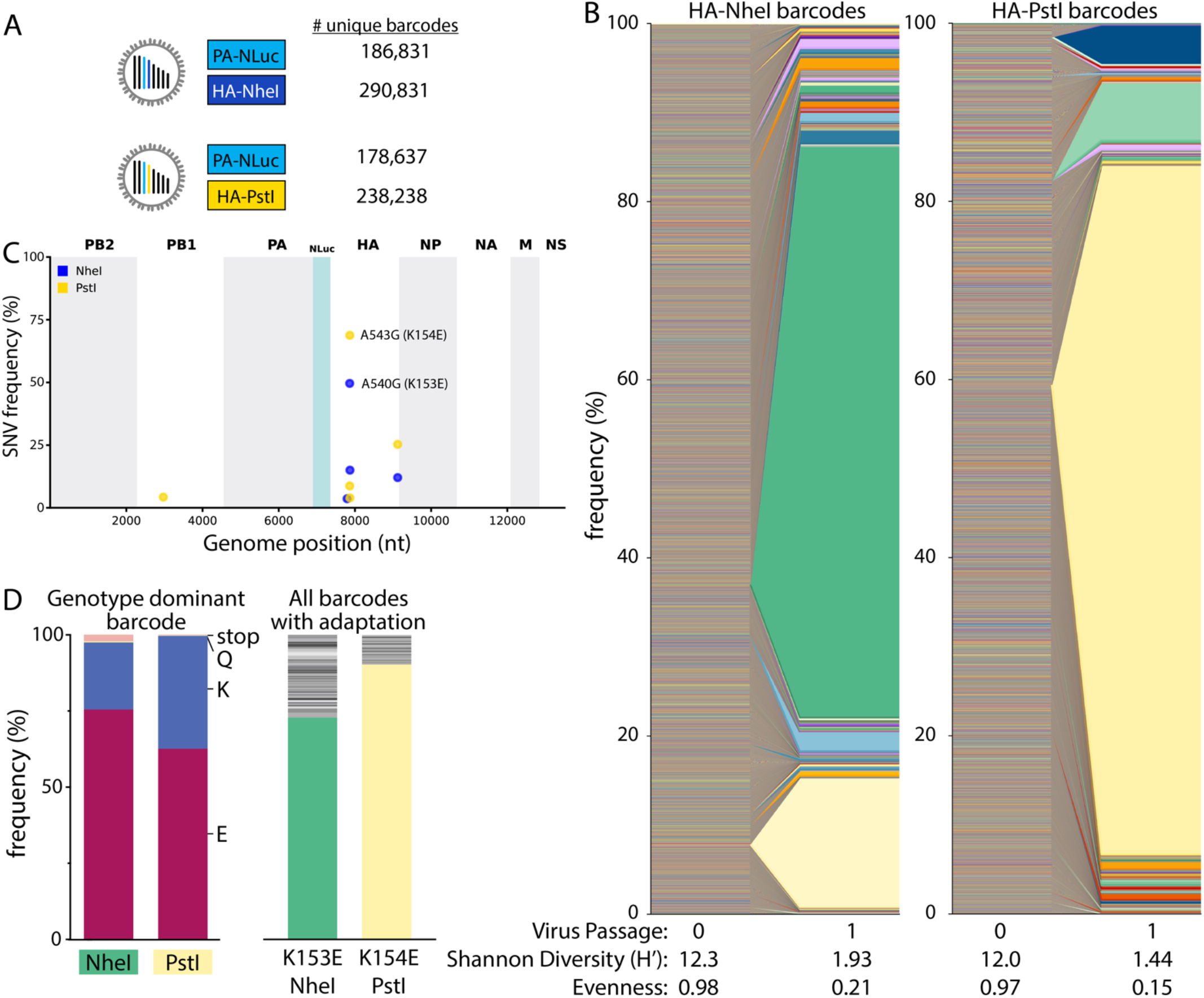
Single viruses seed selective sweeps in *HA*. A) Creation of two dual-barcode libraries with distinct registration marks and uniquely addressable members. B) Frequency of each lineage as a fraction of total population size. Colors indicate unique barcode identities. C) Whole genome sequencing identifies adaptive variants in *HA*. Individual single nucleotide variant (SNV) frequencies are indicated at each nucleotide position in a concatenated IAV genome for each library. D) Long-read sequencing reveals selective sweeps by linking adaptive mutants in HA to single dominant barcodes. The frequencies of mutations coding for the indicated change that are linked to the dominant barcode are shown for both libraries (left). The frequencies of all barcodes linked to the adaptive glutamic acid variant are indicated (right), with the dominant barcode in each library colored as in B.

Changes in population composition can occur through drift or selection. We passaged virus at a large effective population size, minimizing the propensity for drift and raising the possibility that the expanding lineages became dominant because they had acquired a selective advantage. Whole genome sequencing of the P1 stocks identified the single nucleotide variants (SNVs) A540G (numbering based on cRNA) in almost 50% of reads from NheI registration mark viruses and A543G in almost 70% of reads from PstI marked viruses (**Fig 2C**), mirroring the abundance of the dominant barcode. These mutations code for HA K153E and K154E (H1 numbering), respectively. Importantly, these changes in HA had previously been identified as adaptations that provide a growth advantage to CA07 in cell culture^27^. Long-read sequencing showed that the dominant barcode was linked to the adaptive mutations (**Fig 2D, left**). 62-75% of reads containing the dominant barcode also encoded the adaptive mutation. Similarly, the adaptive mutation was primarily linked to the dominant barcode (**Fig 2D, right**). However, up to 27% of reads encoding these adaptive variants were associated with different very low-frequency viral lineages. These data are consistent with a soft selective sweep where *HA* A540G or A543G arose on multiple genetic backgrounds, even though only one lineage ultimately became the most abundant, possibly suggesting clonal interference. These observations demonstrate that our barcoded viruses capture lineage dynamics and selective processes at extremely high resolution.

### Pre-adaptation creates large and diverse viral libraries

To create diverse libraries without tissue-culture-induced skewing, we made new libraries on a “pre-adapted” HA K153E background. The libraries contained at least 57,000 unique members (**Fig 3A**). Multicycle growth curves of HA K153E NheI- or PstI-marked barcoded viruses were indistinguishable (**Fig 3B**). Insertion of the barcode cassette and the *PA* reporter gene slightly reduces titers when compared to HA K153E alone, consistent with prior results^20,28^. The new libraries were highly diverse (Shannon’s Diversity Index (H’) >8.8 for all barcodes) and evenly distributed; the vast majority of lineages were present at low frequency, and no single lineage in our amplified stocks was present at a frequency above 0.6% for HA, or above 1.2% for PA barcodes **(Fig 3C)**. We utilized the HA-NheI stock for extensive quality control of our barcode enumeration pipeline. Libraries were prepared from four independent RNA extractions and sequenced. Lineage frequencies from the four replicates showed strong correlation (**Supp Fig 1A**). Over 94% of all sequence reads were shared across the four replicates (**Supp Fig 1B**). However, when considering barcode identity, only ∼37% of lineages were identified in all replicate runs (**Supp Fig 1C**). This apparent discrepancy is due to variable detection of extremely low-frequency lineages. The frequency of an individual lineage was correlated with the number of replicates in which it was detected (**Supp Fig 1D**). These data highlight the complexity of our viral populations and our ability to reliably detect individual members. Replicate sequencing of the *PA* barcode and the HA-PstI stock produced similarly well-correlated results (**Supp Fig 1E, 2A-B**). No additional SNVs were detected at high frequency in our libraries, while K153E remained fixed **(Fig 3D)**.

**Figure 3.**
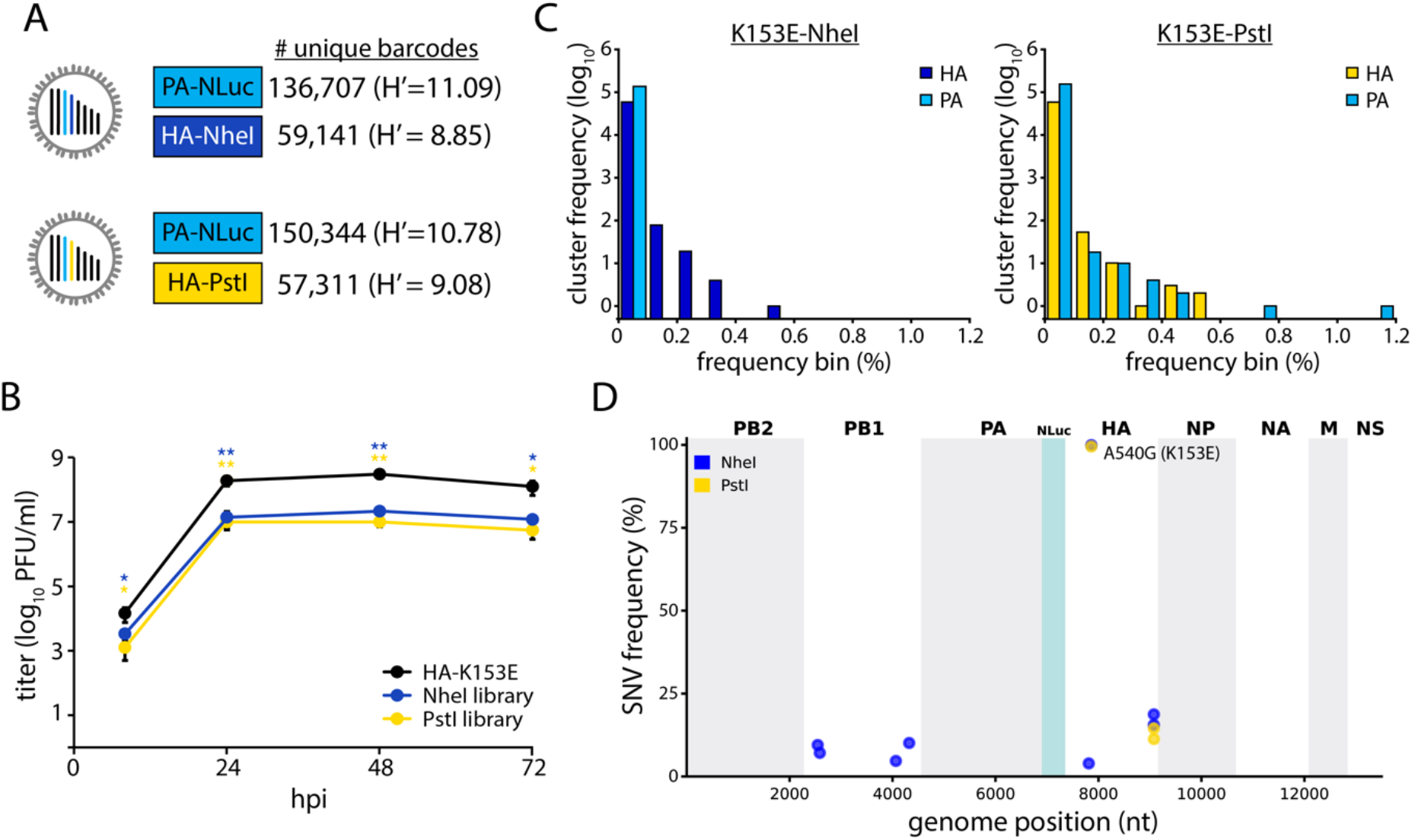
Generation of large and evenly distributed dual-barcoded virus libraries. A) Properties of the A/CA/07/2009 HA-K153E PASTN virus libraries. Data are from a single sequencing replicate; see Supplemental Figs 1-2 for additional analyses. A) Unique barcodes identified in each population. B) Multistep growth curves of dual-barcoded PASTN libraries compared to the parental strain A/CA/07/2009 HA-K153E. Viral titer was measured by plaque assay (n=3 ± sd, ANOVA with Tukey’s post hoc, * = p < 0.05 and ** = p < 0.01 compared to the parental). C) Frequency distribution of unique barcodes on *HA* and *PA* binned in 0.1% increments. D) Whole genome sequencing was performed and SNV frequencies are indicated at each nucleotide position for each library.

### Rich and diverse populations replicate in mice

The dual barcoded virus libraries provide a key opportunity to quantify population dynamics *in vivo*. Mice are frequent models for influenza virus replication, pathogenesis, and immune response^29^. Mice were inoculated with the CA07 PASTN-barcoded virus libraries containing either barcoded HA-K153E populations or a non-barcoded HA-K153E control. Weight loss was similar for all conditions **(Fig 4A)**, and viral titers in the lungs were not significantly different at either 3 or 6 dpi **(Fig 4B)**. Thus, introduction of barcodes onto *HA* does not compromise replication *in vivo*.

**Figure 4.**
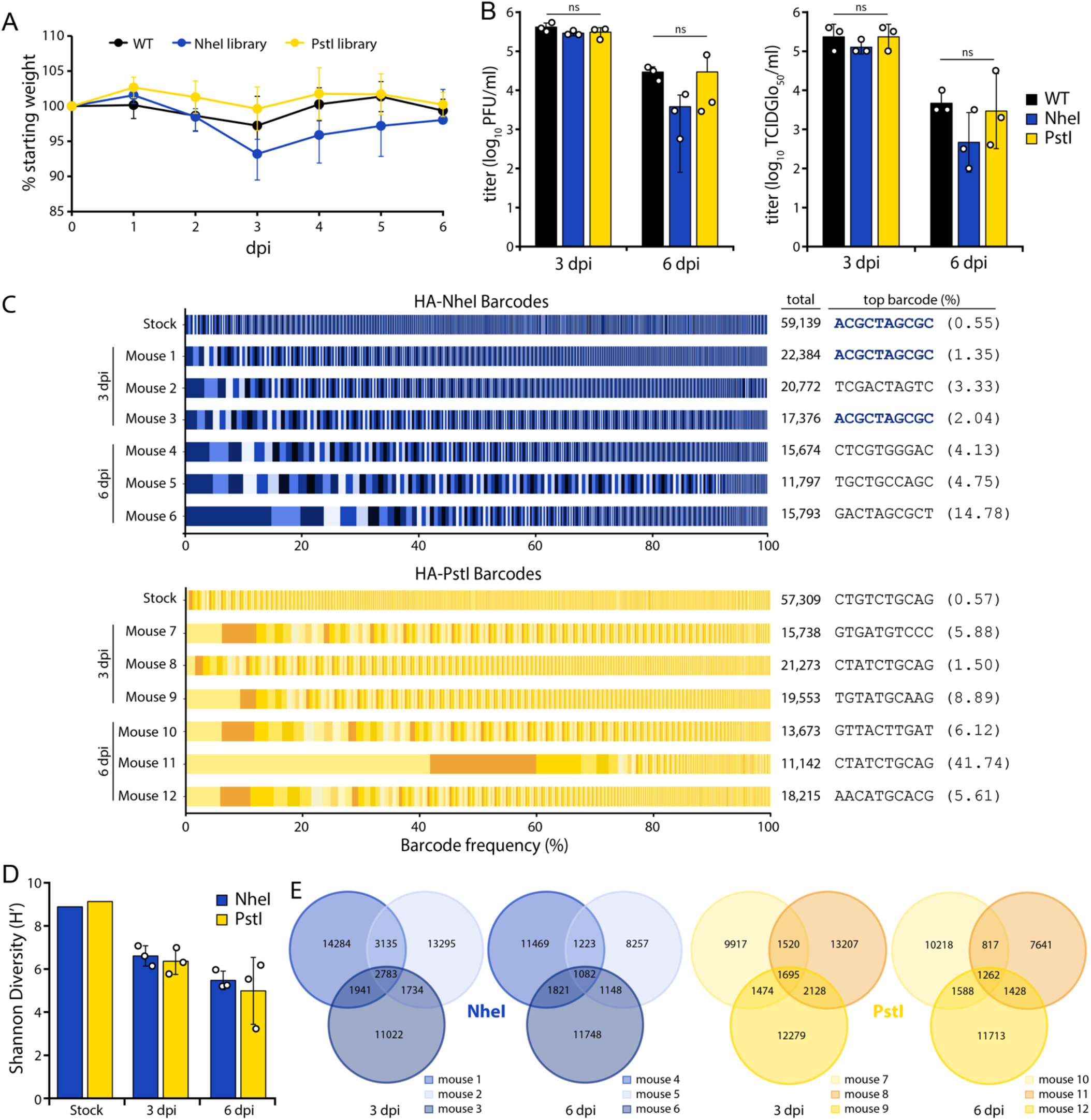
Replication of diverse virus populations in mice. A) Mice were inoculated with 10^5^ TCIDGlo_50_ of virus containing barcoded PASTN with either HA-K153E or barcoded variants and body weights were measured daily. Half of the mice were sacrificed at 3 dpi. Data presented as mean ± sd for n=6 1-3 dpi, and n=3 for 4-6 dpi. B) Viral titers in mouse lungs harvested at 3 and 6 dpi were determined by plaque assay (left) or TCIDGlo_50_/mL (right). Data presented as mean of n=3 ± sd. C) Barcodes in the viral stock and mouse lungs were quantified and the frequency of clustered HA barcodes as a fraction of total population size is indicated. Each color in a series represents an individual barcode cluster. Total number of unique barcode clusters per sample and the most abundant barcode with its frequency are listed at right. D) Shannon’s diversity index for viral populations in the stock and mouse lungs (mean of n=3 ± sd). E) Venn diagrams displaying the number of unique and shared lineages within each mouse for NheI and PstI libraries.

Deep sequencing of mouse lung homogenates revealed that the majority of mice harbored diverse viral lineages **(Fig 4C-D)**. Mouse infections were characterized by a high richness, where approximately one third of lineages present in the stock were detected in the lungs at 3 dpi. The frequency of lineages in the NheI-marked stock was moderately predictive of their abundance in the mouse at 3 dpi (Pearson’s R > 0.57)(**Supp Fig 3A**). For example, the most abundant lineage in the HA-NheI stock was also the most abundant lineage in two mice at 3 dpi (**Fig 4C, Supp Fig 3A)**. However, there are notable exceptions in which low-frequency lineages in the stock rose to high relative abundance in the mouse (**Supp Fig 3A**). In addition, we cannot exclude the possibility that some of our barcodes are derived from residual viruses from the inoculum that did not initiate a productive infection. Mice infected with PstI-marked libraries showed less correlation between the inoculum and lungs at 3dpi (**Supp Fig 3A**). For all viral libraries, the titers decreased from 3 to 6 dpi, as did richness and overall diversity (**Fig 4C-D**). In an extreme example, a single lineage in mouse 11 rose to over 40% prevalence at 6 dpi (**Fig 4C, Supp Fig 3B)**. The correlation between lineage frequency in the stock and the mouse lung was greatly diminished at 6 dpi (**Supp Fig 3)**. The lineage identities are highly heterogeneous among mice, with a small fraction shared across animals, suggesting that barcodes themselves are not under selection *in vivo* **(Fig 4E)**. Together, these data show virus populations replicating in mice approximate the diversity present in the inoculum, as might be expected from a high-dose challenge. Within-host richness decreases as the infection is resolved, with lineages lost as the overall population size decreases.

### Within-host bottlenecks result in compartmentalized replication in the ferret lower respiratory tract

Ferrets are often considered the “gold standard” infection model with lung physiology, sialic acid distribution, pathogenesis, and transmission capacity that are all similar to humans^30,31^. We intranasally inoculated 3 ferrets with the HA-K153E dual barcoded library containing the NheI registration mark. We used a site-specific inoculation strategy in which the inoculum is retained in the upper respiratory tract without unintentional introduction into the trachea or lower respiratory tract^32^, allowing us to track the natural movement of virus. Ferrets were monitored daily for signs of infection with nasal washes obtained 1, 3 and 5dpi. Ferrets exhibited slight weight loss over the course of infection **(Fig 5A)**, consistent with prior work^28^. Similarly, we detected high viral titers in nasal washes 1 dpi that declined over time **(Fig 5B)**.

**Figure 5.**
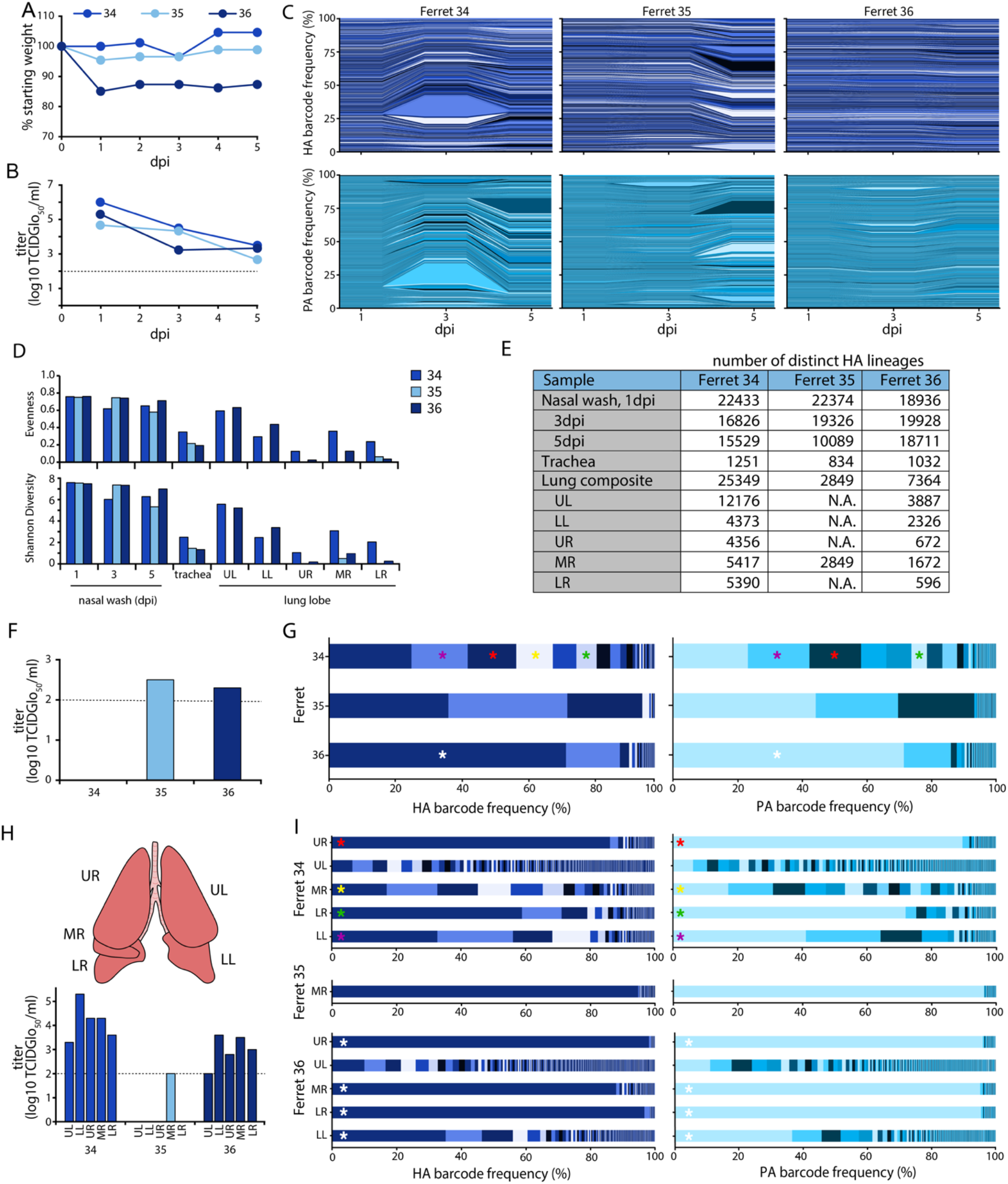
Population diversity is reduced when influenza virus moves from the upper to the lower respiratory tract in ferrets. A) Ferrets were inoculated with a site-specific intranasal dose of 10^5^ PFU of dual-barcoded A/CA/07/2009 HA-K153E PASTN virus containing the NheI registration mark. Ferret weight was monitored daily. B) Viral titer in ferret nasal washes were determined by TCIDGlo_50_. C) Changes in frequency for HA (top) and PA (bottom) barcodes present in nasal wash samples over the course of infection. D) Shannon’s diversity index and evenness for the indicated viral populations. E) The richness of each tissue is indicated by the number of distinct barcodes in each sample. N.A. = not attempted. F-G) Virus was recovered from the trachea at 5 dpi. F) Viral titers were measured by TCIDGlo_50_ and G) the frequency of barcodes was determined. Distinct dominant barcodes are identified by colored asterisks. H) Viral populations in individual lung lobes were titered by TCIDGlo_50_ and I) the frequency of barcodes was determined. Lung lobes: upper left (UL), lower left (LL), upper right (UR), middle right (MR), lower right (LR). Distinct dominant barcodes are identified by colored asterisks, matching those in G) where appropriate. The limit of detection for viral titer assays is indicated by a dashed line in B, F, and H.

Sequencing revealed rich and complex viral populations in the nasal washes of ferrets, consistent with a high dose inoculation in this compartment (**Fig 5C-E**). Frequency trajectory plots showed heterogeneous and well mixed viral populations undergoing little, if any, selection or bottlenecks in the upper respiratory tract (**Fig 5C**). This is consistent with a highly diverse population with generally even lineage distribution, reflected by a high Shannon’s diversity index and evenness (**Fig 5D**). Approximately 19,000-22,400 unique lineages were detected in each nasal wash 1 dpi, and richness remained high over time, with at least 16,800 lineages at 3 dpi and 10,000 lineages at 5 dpi **(Fig 5E)**. Sequencing of the barcode on *PA* revealed remarkably similar lineage dynamics. The fact that frequencies of *HA* and *PA* barcodes moved in parallel is perhaps surprising given that these are unlinked genes and that influenza viruses can undergoes frequent reassortment^33^.

Intranasally inoculated viruses spread throughout the respiratory tract by 5 dpi. Low levels of infectious virus were present in the trachea of ferrets 35 and 36, whereas deep sequencing detected IAV genetic material in all trachea samples (**Fig 5F-G)**. Compared to the nasal wash, population richness dropped significantly, with 1250 or fewer lineages present in the trachea. Lineage distribution differed in the trachea of each animal, ranging from a more diverse population in ferret 34 to a largely homogenous population in ferret 36, in which a single lineage accounted for 71% of the population **(Fig 5G)**. Our site-specific inoculation requires virus replication in the upper respiratory tract prior to movement into the trachea or lower respiratory tract. We therefore used lineages present in nasal washes at 3 dpi as a comparator for populations in the trachea and lungs at 5 dpi. Lineage frequency is poorly correlated between nasal washes and the trachea (**Supp Fig 4A**). Migration into the trachea is associated with a drastic reduction in richness, a poor correlation with the source, and skewed distribution of the resultant population, indicating that viral population bottlenecks between compartments and founder effects may play a role during the seeding of the trachea from the upper respiratory tract.

Virus also spread to the lungs of infected animals **(Fig 5H)**. Moderate titers were detected in all five lung lobes in ferret 34, even though infectious viral titers in the trachea were below the limit of detection for this animal. Infectious virus was also detected in all lung lobes for ferret 36, but only the middle right lobe for ferret 35. Thus, while virus must traverse the trachea to access the lungs, the presence of virus in the trachea 5 dpi was not predictive of the extent of spread in the lungs. Moreover, populations in lung lobes had higher lineage richness than that detected in the trachea at 5 dpi; the vast majority of lineages in the lung were not detected in the trachea (**Supp Fig 4B**). This raises the possibility that population richness in the trachea at 5 dpi shrank significantly from the populations at earlier time points that may have seeded infection in the lung, or that virus can transit through the trachea to directly inoculate the lung.

Lineage analysis revealed heterogeneous populations of barcodes in each of the distinct lobes (**Fig 5I)**. Over 25,000 lineages were detected across the five lobes in ferret 34. However, each lung lobe of ferret 34 had a different dominant barcode sequence, and when this same barcodes was detected in other lung lobes its frequency varied. The only infected lung lobe of ferret 35 had very low viral titers and was dominated by a single lineage reaching ∼95% abundance (**Fig 5H-I**). Ferret 36 yielded another outcome, where the same lineage was dominant at a frequency of 35-98% in each lobe except for the upper left, which maintained a richer and more diverse population. For all animals, individual lobes showed reduced diversity and evenness compared to the virus population in the upper respiratory tract (**Fig 5D**). In the two animals in which virus was detected in all lobes, the upper left lobe consistently had the highest richness, diversity, and evenness. These data suggest that anatomical features associated with each lobe, such as tracheal bifurcation patterns or bronchus size^32^, may affect patterns of virus establishment and population size. The differences in composition of the populations in each lung lobe was determined. Bray-Curtis dissimilarity assessment revealed highly compartmentalized replication, in which each lung behaved as a distinct anatomical “island” with a unique population composition (**Fig 6A**). Four lobes in ferret 36 showed lower dissimilarity, as they were dominated by the same lineage, yet the non-dominant lineages still contributed unique populations to each lobe. Only 167 lineages were common to all lobes of ferret 34, and only 42 in ferret 36 (**Fig 6B**).

**Figure 6.**
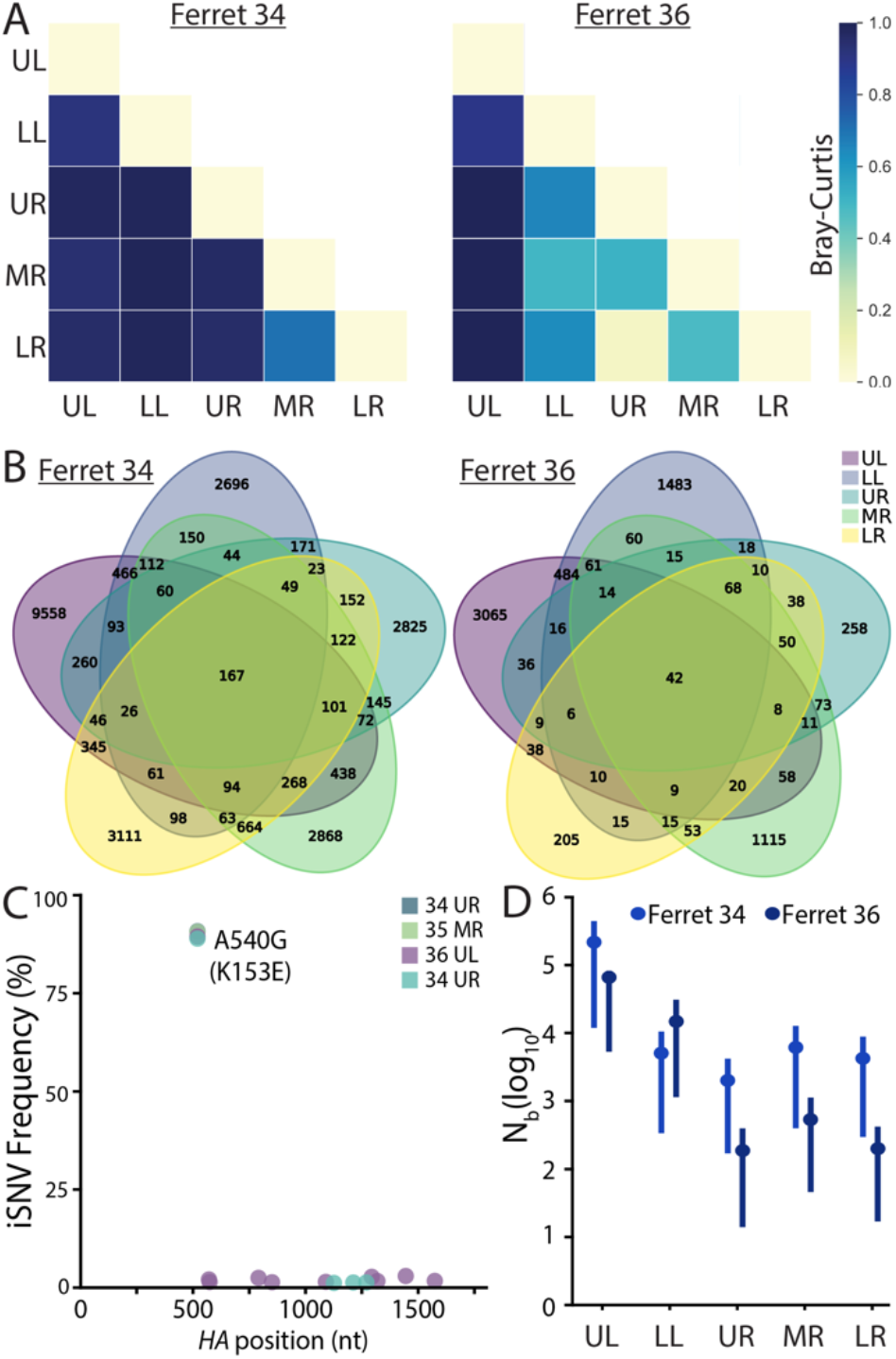
Lung lineage diversity and population bottlenecks are dominated by stochastic pressures. A) Pair-wise Bray-Curtis dissimilarity for lung lobes within single animals. B) The number of barcode lineages common and unique to lung lobes are illustrated. C) Whole-HA sequencing of virus in lung homogenates identified multiple iSNVs, but not rose above the 3% threshold set for accurate estimation of iSNV frequency. D) Transmission bottlenecks (*N*_*b*_) from nasal wash 3 dpi to each lung lobe at 5 dpi were calculated by maximum likelihood estimation and plotted ± 95% confidence intervals.

The drastic reduction in richness and diversity between the upper respiratory tract and the lung lobes suggested populations were subject to selection or bottlenecks. *HA* sequencing from select lung lobes for each animal revealed that no intrahost SNVs (iSNVs) surpassed the 3% threshold set for accurate estimate of iSNV frequency, excluding the possibility that barcode frequency changes were driven by selection of a linked adaptive variant (**Fig 6C**). We therefore further assessed the possibility of viral population bottlenecks in driving the observed reduction in richness and diversity. To this end, we developed a simple mutlinomial model to estimate bottleneck sizes (*N*_*b*_) as virus transmits from the nasal wash at 3 dpi to lung lobes at 5 dpi (**Fig 6D**). Specifically, this model estimated *N*_*b*_ using data on the lineages and their frequencies in the 3 dpi nasal wash and on the number of lineages observed in a focal 5 dpi lung lobe. The model yielded maximum likelihood estimates of *N*_*b*_ of ∼66,000 (ferret 36) and ∼217,000 (ferret 34) virions for the upper left lobe, and lower estimates for the other lobes (**Fig 6D**), with maximum likelihood estimates spanning between ∼200 and ∼15,000 virions. The larger bottleneck size between the nasal wash and the upper left lobe compared to that for the other lung lobes was consistent with higher levels of viral lineage diversity detected in the upper left lobe compared to the others. Nonetheless, the bottleneck size estimates still appeared unexpectedly large to us, given the drastic reductions in richness and diversity observed in Fig 5D. We therefore forward simulated mock transmission events from the nasal wash using our estimated bottleneck sizes under the assumption that virions were transmitted from the nasal wash to a focal lung lobe at a single timepoint. These simulations predicted a high degree of similarity between lineage frequencies in the viral populations sampled in the nasal wash and lung lobes (**Supp Fig 5**). They further predicted levels of viral genetic diversity and richness in the lung lobes that were significantly higher than those observed (**Fig 5D,I**). These initially seemingly inconsistent results could be parsimoniously explained under a model where many virions are transmitted between the nasal wash and each of the lung lobes over the course of infection, but at multiple timepoints rather than at a single one.

Considering the lung as a whole for ferrets 34 and 36, a substantial number of lineages were shared between the nasal wash and lung (**Fig 7A**). But, this appeared to be largely driven by the rich and diverse population in the upper left lobe, as this overlap was largely lost when lobes were considered individually. Many of the dominant lineages in a lung lobe were poorly represented in the nasal wash, such as that in the upper right lobe of ferret 34 and the dominant lineage shared in 4 lobes of ferret 36 (**Fig 7B, C, Supp Fig 6A, C**). Lineage enrichment in the lung samples compared to nasal washes revealed many lineages that were unique to nasal washes or lung lobes (**Sup Fig 6A-C**). These data show that treating the lung as a whole can lead to very different and misleading population structures, highlighting the importance of assessing each lobe individually.

**Figure 7.**
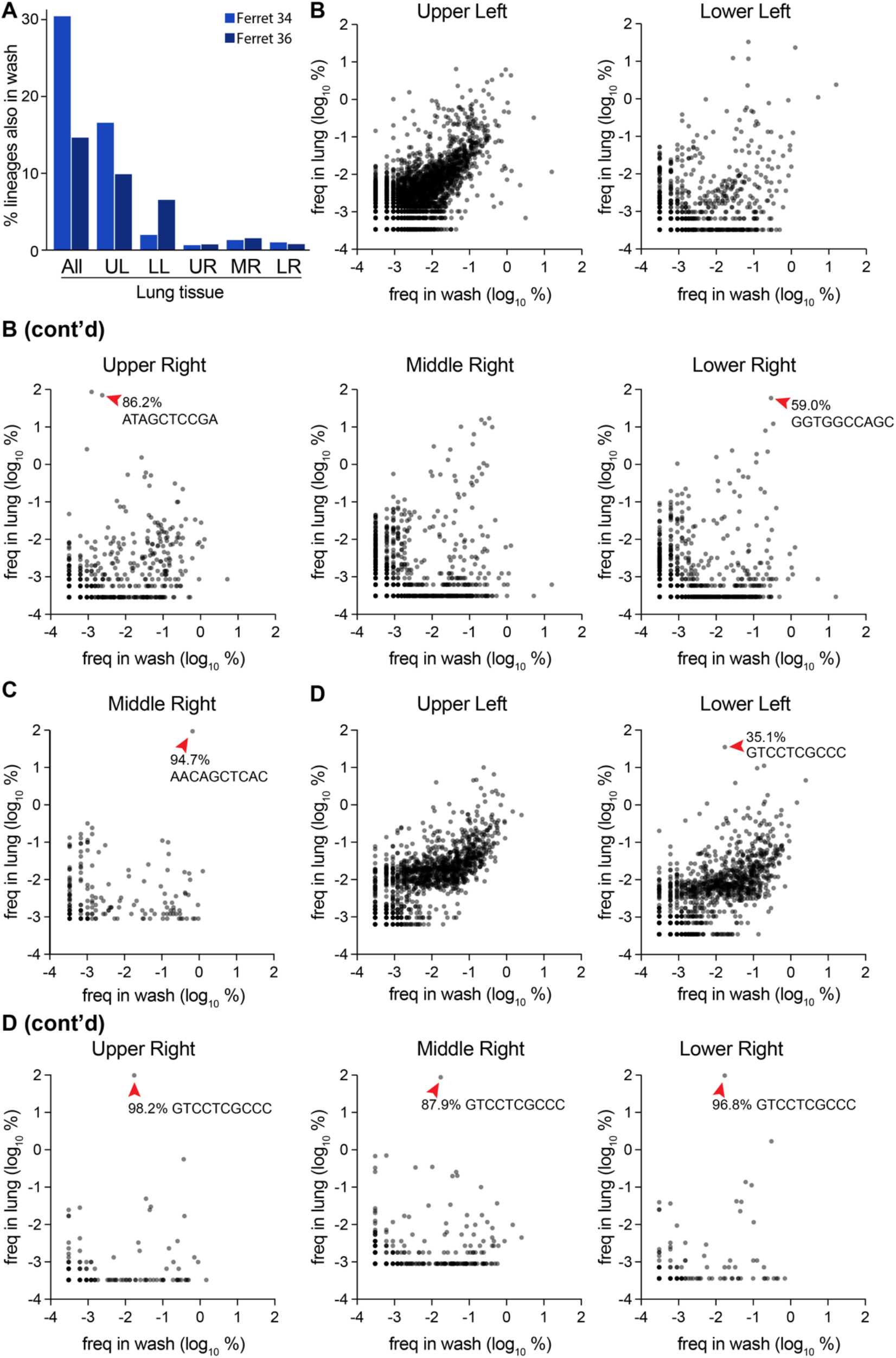
Compartmentalized infections establish replication islands within the lung. A) Overlap between barcodes in the nasal wash 3 dpi and those present in distinct lobes, or composite data for all barcodes in the lung. B-D) The frequency of barcodes within the nasal wash are compared to frequencies within individual lobes for B) ferret 34, C) ferret 35 and D) ferret 36. Red arrowheads highlight dominant barcodes with frequency > 30%, which for ferret 36 is the same barcode that dominated in multiple lobes of the lung. Note that barcodes unique to the nasal wash or lung are not plotted here and can be found in Supplemental Figure 6.

The same lineage dominated in four lobes of the lung for ferret 36, whereas underlying diversity and a lack of overlap in each population still suggests each lobe is seeded by distinct transmission events (**Fig 5I, 6B 7D)**. In this animal, enrichment for the dominating lineage appeared to occur in the trachea (**Fig 5G, Supp Fig 4A, 7A**). This may be a general trend, as high-frequency lineages in the trachea were often over-represented in the lobes (**Supp Fig 7A-B**). Combined, the large differences in lineage identity and frequency show that each lobe is independently seeded, with little mixing between compartments. Moreover, comparisons across compartments suggest each lobe presents a bottleneck and that genetic drift repeatedly reshapes the population as virus traverses the respiratory tract.

## DISCUSSION

A quantitative understanding of population dynamics is crucial for determining how evolutionary forces shape viral populations within and between hosts. Error-prone replication by influenza virus generates genetic diversity within an infected host. However, the full extent of that diversity does not survive transmission to a new host. Transmission events between hosts involve stringent bottlenecks, with only 2-6 viral genomes founding the next infection during aerosol transmission^8,10,34^. Here we show that influenza virus also faces multiple bottlenecks within a host as it seeds different compartments. Rich and diverse populations in the upper respiratory tract were stochastically sampled as virus transited into the lower respiratory tract, introducing strong founder effects that skewed the resultant population. Our data suggest a scenario in which repeated within-host bottlenecks severely reduce diversity, richness, and evenness, resulting in distinct, compartmentalized “island populations” in each lobe of the lung.

Our results show that influenza virus evolution within a host involves multiple bottlenecks, tempering the impact of positive selection and likely creating additional barriers for host-adaption and onward transmission. This could involve multiple physical bottlenecks as the virus sequentially infects distinct tissues, repeated exposure to the same bottleneck if transmission to a new tissue occurs more than once, or a combination of the two. And while we considered entire lung lobes, it remains possible that focal infections within tissues may further subdivide viral populations. In all scenarios, bottlenecks decrease the richness and overall population size and are expected to increase the strength of genetic drift^35^. These processes combine to constrain the diversity and evolution during influenza virus infection, which slows the fixation of adaptive variants within a host.

Immune pressure positively selects antigenically advanced influenza virus variants on the global scale^1^. Yet, these same variants rarely emerge in an acutely infected host, where infections are dominated by purifying selection and low intra-host diversity^4,5,7,8,36^. The apparent disconnect between influenza virus evolution on the global versus individual scale is incompletely understood. Our findings reveal that intrahost bottlenecks are major contributors to the limited evolution detected in an infected individual. This may partially explain why highly pathogenic avian H5N1 strains have not yet acquired the ability to transmit within people. H5N1 strains infect the lower respiratory tract in humans, where the preferred α2,3-sialic acid receptors are more abundant^15,16^. We saw little evidence for mixing between compartments in the lower respiratory tract. Thus, while only a few mutations are needed to confer airborne transmission to H5N1 strains in experimental settings, the spatial structure and compartmentalization of infection in humans may impose bottlenecks that prevent fixation of variants and migration to the upper respiratory tract, let alone transmission to a new host^37,38^.

Intrahost bottlenecks are common features during viral infections. Physical barriers establish bottlenecks as enteroviruses escape the gut in mouse models^39,40^. Poliovirus then faces another bottleneck related to IFN responses that restrict diversity as virus invades the central nervous system^39,41,42^. Our data do not reveal whether the bottlenecks we describe for influenza virus are due to physical barriers, innate immune response, or other factors. Similar forces shape evolution in arthropod vectors. Repeated bottlenecks winnow diversity as virus moves between different tissues within mosquitoes for West Nile virus, Venezuelan equine encephalitis virus and Zika virus^35,43,44^. Error-prone replication then repopulates diversity in the new sites of replication. Repeated bottlenecks, such as those we have identified during intra-host dissemination, decrease diversity and may reduce fitness. Yet, they also act to purge deleterious mutations and facilitate escape from local fitness maxima and exploration of additional evolutionary space. The combined effects of these bottlenecks shape the kinetics of intra-host influenza virus evolution.

In summary, we demonstrate the existence of multiple bottlenecks during dissemination of influenza virus throughout the respiratory tract. We posit that these bottlenecks contribute to the limited impact of positive selection on intrahost evolution. Moreover, they provide additional barriers to initial cross-species transmissions and sustained transmission once a virus spills over into a new host. Coupled with the stringent bottlenecks that occur during inter-host transmission, our results help explain the stochastic nature of influenza virus evolution at the local scale.

## METHODS

### Cells

MDCK cells (ATCC), MDCK-SIAT1-TMPRSS2 cells^45^ and 293T cells (ATCC) were maintained in Dulbecco’s modified Eagle medium (DMEM) supplemented with 10% heat-inactivated fetal bovine serum (Atlanta Biologicals), 100 μg/mL streptomycin, and 100 U/mL penicillin at 37°C and 5% CO_2_.

### Generation of pHW2000-all-ΔPA-ΔHA A/California/07/2009

Bidirectional cassettes for CA07 PB2, PB1, NP, NA, M, and NS gene segments were sequentially amplified and inserted into a pHW2000 vector via Gibson assembly. The resulting 17.3 kb plasmid was sequence verified. Annotated plasmids sequences can be found at https://github.com/mehlelab/barcoded_flu_analysis

### Generation of barcoded PA

Barcodes were inserted into the previously described PASTN rescue construct that expresses a polyprotein encoding PA and Nanoluciferase separated by the 2A cleavage site^20,46^. The barcode was originally synthesized (IDT) as a single-stranded DNA oligo containing ten random nucleotides flanked by sequence to enable cloning immediately downstream of the Nanoluciferase open reading frame (Supplementary Table 1). The oligo was amplified in 25 individual low-cycle PCRs using Q5 polymerase (NEB).The amplicons were pooled and gel purified prior to cloning into a modified PASTN plasmid using 50 individual ligation reactions. Ligations were transformed into Mach1 competent *E. coli* (Thermo Fisher), plated on LB agar supplemented with 200 μg/mL ampicillin, and grown overnight at 37°C to yield 220,000 transformants, indicative of a theoretical maximum library size. Colonies were scraped off plates, collected in 750 mL liquid LB with ampicillin, and grown for 3 hr at 37°C. DNA was then purified using the Zymo Midiprep kit to create the plasmid library. Plasmid stocks were deep sequenced to determine library size and diversity.

### Generation of barcoded HA

*HA* from A/California/07/2009 was cloned into the pHW2000 rescue plasmid. Silent mutations were introduced into the final 25 amino acids of the open reading frame and 80 nt from the 3’ end of the open reading frame were repeated downstream to recreate a contiguous packaging signal^21^. Initial libraries utilized the native HA sequence, whereas subsequent libraries included the tissue-culture adaptive K153E (nt A540G in cRNA) mutation^27^. A single-stranded oligo (IDT) was synthesized containing a randomized 10-nt barcode as well as a 6-nt registration mark, either GCTAGC (NheI) or CTGCAG (PstI) (Supplementary Table 1). This registration mark is a conserved and identifiable region for all the individuals within a given library. Barcodes were amplified in 50 low-cycle PCRs and cloned following the same strategy as for PA libraries. Approximately 60,000 transformants were obtained with the NheI registration mark, and 60,000 for the PstI registration mark. Plasmid stocks were deep sequenced to determine library size and diversity.

### Rescue of dual-barcoded virus libraries

Barcoded CA07 virus libraries were rescued via reverse genetics using the pHW2000-all-ΔPA-ΔHA and PA and HA barcoded plasmids described above. Briefly, 293T were forward transfected with 2.7 μg pHW2000-all-ΔPA-ΔHA, 450 ng pHW2000-PASTN-barcode library, 450 ng pHW2000-HA-barcode library (NheI or PstI variant), and 400 ng pHAGE2–EF1ɑInt–TMPRSS2–IRES–mCherry-W^45^ in a 6-well format. Plasmids were combined with 200 μL jetPRIME Buffer and 8 μL jetPRIME reagent (Polyplus) per well. 120 independent transfections were performed per viral library. 24 hr post-transfection, media was removed and cultures were overlaid with MDCK-SIAT1-TMPRSS2 cells in OptiVGM (OptiMEM supplemented with 0.3% bovine serum albumin, 100 μg/mL calcium chloride, 100 μg/mL streptomycin, and 100 U/mL penicillin). Rescue viruses were harvested 48-72 hr later and pooled based on their registration mark. Viruses were amplified on 20, 15 cm dishes of MDCK-SIAT1-TMPRSS2 cells in OptiVGM for 66 hr. Viruses were pooled based on their registration mark and cellular debris was removed by centrifugation. Viral titers were determined by plaque assay and TCID50Glo assays on MDCK and MDCK-SIAT-TMPRSS2 cells^47^.

### Library preparation for barcoded amplicons

Viral RNA was extracted from all samples using the Maxwell RSC Viral Total Nucleic Acid Purification Kit (Promega) according to the manufacturer’s instructions. RNA was subjected to DNAse treatment using the TURBO DNAse (Invitrogen), and reverse transcribed in 20µl using the SuperScript IV VILO master mix (Invitrogen) with PA and HA gene segment-specific primers (Supplementary Table 1). DNA amplicons for HA and PA gene segments with partial sequencing adapters were generated via PCR amplification of cDNA using the Phusion High-Fidelity DNA Polymerase (New England BioLabs) and gene segment specific primers (Supplementary Table 1). To minimize technical bottlenecks during library preparation, reverse transcription and the first PCR amplification for all samples were performed in triplicate and pooled together prior to DNA purification. PCR products from mouse samples were gel purified whereas PCR products from ferret samples were purified with paramagnetic beads using the AMPure XP for PCR Purification kit (Beckman Coulter). Purified PCR products were used in a second PCR reaction for incorporating sample-specific 5’-end indexes and additional Illumina sequencing adapters (Supplementary Table 2). Final PCR products were gel purified and individual DNA concentrations were determined with the Qubit dsDNA High Sensitivity Assay Kit on the Qubit Fluorometer (Invitrogen). Samples were quality controlled using the Bioanalyzer High Sensitivity DNA Analysis Kit and the Agilent 2100 Bioanalyzer (Agilent). All samples were prepared and sequenced in technical replicate. Detailed protocols and code available at https://github.com/mehlelab/barcoded_flu_analysis.

### Library preparation for whole-genome sequencing

Library preparation was similar to our prior approaches^48^. Briefly, viral RNA was extracted from all samples using the Maxwell RSC Viral Total Nucleic Acid Purification Kit (Promega) according to the manufacturer’s instructions. RNA were subjected to DNAse treatment using the TURBO DNAse (Invitrogen), and reverse transcribed in 20µl using the SuperScript IV VILO master mix (Invitrogen) with the Uni12 primer (Supplementary Table 1) that targets conserved ends of all gene segments. Segments were amplified by PCR with gene-specific primers (Supplementary Table 1), gel purified, and DNA concentrations were determined using the Qubit dsDNA High Sensitivity Assay Kit on the Qubit Fluorometer (Invitrogen). 1 ng of each segment was pooled and used as input for the Nextera DNA Library Prep kit where samples were tagmented and indexed according to the manufacturer’s instructions. Tagmented and amplified products were purified with AMPure XP paramagnetic beads for PCR Purification kit (Beckman Coulter) in two consecutive steps (0.5x and 0.7x) and were quantified using Qubit dsDNA high-sensitivity kit (Invitrogen, USA). Sample quality control was performed using the Bioanalyzer High Sensitivity DNA Analysis Kit in the Agilent 2100 Bioanalyzer (Agilent). All samples were prepared and sequenced in technical replicate. Detailed protocols can be found in https://github.com/haddocksoto/bcflu_protocols.

### Deep sequencing

Amplicon and whole-genome libraries were sequenced on the Illumina MiSeq system using the MiSeq Reagent Kit v2-500 and v3-600, respectively (Illumina). Amplicon and whole-genome samples that passed quality control were pooled in a 4nM library with nuclease-free water. 5 μl of the 4nM library pool was denatured with 5 μl 0.2N of NaOH and diluted using the HT1 Hybridization Buffer (Illumina) to a concentration of 8 pM for amplicon samples and 10 pm for whole-genome samples. A PhiX library was prepared similarly, and added at 30% of the input for amplicon sequencing and 1% for whole-genome libraries. Samples were loaded on the respective MiSeq cartridge and paired-end sequencing reads were generated (Illumina).

### Sequencing data analysis for barcoded amplicons

We generated a custom bioinformatic pipeline to process raw FASTQ files and quantify barcode frequencies (https://github.com/mehlelab/barcoded_flu_analysis). Briefly, raw FASTQ paired reads were demultiplexed, merged and aligned to a custom amplicon reference containing barcode and registration mark regions as strings of N’s (BBMap Tools v38.87). Reads with lengths the same as the average insert size were aligned, sorted and indexed with Samtools (v1.11, htslib v1.11). BAM files with aligned reads were processed and trimmed to the region containing the registration mark and/or the barcode sequence using command line tools (Seqtk v1.3, Bash v3.2.57).

Sequencing of the invariant registration mark was used to benchmark fidelity of our sequencing. Over 99.1% of reads were perfect matches, where most differences were the result of a single nucleotide change from the expected sequence. Therefore, to correct for amplification and/or sequencing errors that may inflate the number of unique barcodes, we used UMI-tools (v1.1.1) to generate consensus barcode clusters with read counts via the adjacency network-based clusterer method^49^. Parental barcodes and their apparent mutational offspring were clustered prior to enumeration and cluster frequencies were generated and visualized via a custom Python pipeline (Python v3.8.5, Pandas v1.1.3, Matplotlib v3.3.2, Numpy v1.19.2) or in Prism 9. Manual inspection of sequences in the raw FASTQ files confirmed these as *bona fide* barcodes and not the product of misalignment.

### Sequencing data analysis for whole gene segments

We generated a custom bioinformatic pipeline to process raw FASTQ files and determine single nucleotide variants (SNV) from our barcoded viral samples (https://github.com/mehlelab/barcoded_flu_analysis). Briefly, raw FASTQ paired reads were demultiplexed and merged with a minimum overlap region of 30 nucleotides using bbmerge (BBMap Tools v38.87). We mapped reads to the full IAV genome using a Burrows-Wheeler alignment (BWA v.0.7.17). Using Samtools (v1.11, htslib v1.11), we sorted our aligned reads and called variants using LoFreq (v2.1.5) with a minimum coverage of 500 reads, base call quality of at least 30 and a frequency exceeding 0.03 (3%). Our reference IAV genome included barcode insertions in *PA* and *HA*, registration marks on *HA* as a strings of 10 N’s, NanoLuc inserted in *PA*, and repeated packaging signals. SNVs were annotated using SnpEff (v5.0e) to determine the impact of each variant on the amino acid sequence, and the resulting variant call format (VCF) files were manipulated using bcftools (v1.11) to transform into user-defined formats. Plots were generated using custom bioinformatic pipelines in Python language (Python v3.8.5, Pandas v1.1.3, Matplotlib v3.3.2, Numpy v1.19.2).

### Long-read sequencing of *HA* and analysis

Viral RNA was made compatible for sequencing on an Oxford Nanopore Technologies instrument using the 1D PCR Barcoding Kit. Briefly, viral RNA was extracted using the Maxwell RSC Viral Total Nucleic Acid Purification Kit (Promega), subjected to DNAse treatment using TURBO DNAse (Invitrogen), and reverse transcribed and amplified using the SuperScript IV One-Step RT-PCR system with HA-specific primers. Amplified DNA was gel purified using the QIAquick Gel Extraction Kit (Qiagen) and quantified using the Qubit dsDNA High Sensitivity Assay Kit on the Qubit Fluorometer (Invitrogen). Normalized samples were made compatible for long-read sequencing using the 1D Native Barcoding ONT Kit (SQK-LSK-109), following the manufacturer protocol. Libraries were loaded onto a flow cell and run on the ONT GridION machine. Bases were called in real time using the ONT software package Guppy 3.2.6. Minimap2 was used to map reads to the influenza virus *HA* segment and discard low-quality reads^50^. The Sam2Tsv module from Jvarkit^51^ was used to convert the sam to a tsv file so Pandas could be used to extract the bases at position 540 (NheI library) or 543 (PstI library) and the barcode sequence. Results were visualized with Prism 9.

### Sequencing data availability

All sequencing files have been deposited as BioProjects PRJNA746307, PRJNA746319, and PRJNA746307 with details and SRA accessions in Supplementary Table 3.

### Growth kinetics

Triplicate dishes of confluent MDCK-SIAT1-TMPRSS2 cells were inoculated at an MOI of 0.01 with virus diluted in OptiVGM. Virus was adsorbed for 1 hr at 37°C, removed from cells, and replaced with fresh OptiVGM. Virus was sampled over time and titered by plaque assay on MDCK-SIAT1-TMPRSS2 cells.

### Mouse infections

All mouse experiments were approved by the University of Wisconsin Madison Institutional Animal Care and Use Committee. 18 9-week old female BALB/c mice (Charles River Labs) were randomly divided into groups of 6 to receive either WT HA, HA-K153E-NheI-bc, or HA-K153E-PstI-bc virus. All viruses contained barcoded PASTN. Mice were inoculated intra-nasally with 10^5^ TCIDGlo50 of virus in 35 μL media. Mice were weighed daily and monitored for clinical signs of infection. 3 animals from each group were sacrificed at 3 dpi and the remainder at 6 dpi. Lungs were removed, dounce homogenized in 1x DPBS, and clarified at 2000 x g for 5 min. The clarified homogenate was titered via plaque assay and TCIDGlo50 assay on MDCK-SIAT1-TMPRSS2 cells. Viral RNA was recovered from homogenate and sequenced as described above.

### Ferret infections

All ferret experiments were approved by the St Jude Children’s Research Hospital Animal Care and Use Committee. 12-week old male ferrets (Triple F Farms, Sayre, PA) confirmed to be seronegative for influenza virus were housed individually. Each animal was infected intranasally following previous approaches that ensure site-specific inoculation of the upper respiratory tract^32^. Animals were inoculated with 5×10^5^ PFU HA-K153E-NheI-bc with barcoded PASTN diluted in PBS (Corning) containing 100 U/mL penicillin and 100 ug/mL streptomycin (Corning) in a total volume of 500 μl. Nasal washes were collected at 1, 3, and 5 dpi. Briefly, animals were anesthetized with 0.25 mL ketamine and nasal washes collected by administering 1 mL PBS/pen-strep dropwise onto the nostrils. Animals were sacrificed at 5 dpi and trachea and separated lung lobes were removed and frozen prior to processing. Tissue was homogenized in 10% (w/v) L-15 media and using an OMNI TH220-PCRD homogenizer and clarified at 500 x g for 10 min to remove cell debris. Viral titers in homogenates and nasal washes were determined by TCIDGlo50 assay. Viral RNA was recovered from homogenate and sequenced as described above.

### Statistical analysis

Viral titers are presented as the mean of n=3 and significance was tested with a two-way ANOVA with Tukey’s post hoc analysis. Replicate sequencing runs were analyzed with a Pearson’s and Spearman’s correlation coefficient (Prism 9). Population diversity and evenness were assessed by measuring Shannon’s diversity (H’)^26^ as follows:

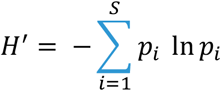

Where *S* is the total number of barcodes detected (richness) and *p*_*i*_ is the frequency of the *i*-th barcode in that sample. Population evenness is bounded at 0 and 1 and defined as the actual barcode diversity divided by the maximum possible diverstiy (H’_max_) for the sample^25^:

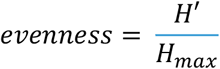

Bray-Curtis dissimilarity (BC_ij_) was used to assess compositions and compare lung lobes^52^, defined as:

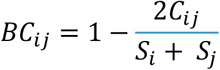

where C_*ij*_ is the sum of the lesser values for only barcodes found in both lobes. *S*_*i*_ and *S*_*j*_ are the total number of barcode reads detected in either lobe.

### Multinomial model of bottleneck size

While several methods currently exist that estimate viral transmission bottleneck sizes between a donor sample and a recipient sample, none of these methods are appropriate for estimating Nb from the barcode lineage frequencies available from this study. The beta-binomial approach outlined in^53^ assumes that that each locus is biallelic and unlinked to other loci. A more recent approach developed by^54^ allows for linked loci by reconstructing haplotypes. However, the number of haplotypes that can be considered in their approach is very low relative to the number of barcode lineages observed in this study. We thus developed a simple statistical approach to estimate Nb that allows for a large number of observed haplotypes (here, barcode lineages). The approach is as follows:

1. Identify the barcode lineages observed in the donor sample and calculate their frequencies
2. Calculate the overall number of barcode lineages present in the recipient sample that are also found in the donor sample
3. Over a range of Nb values, for each Nb do as follows:
  a. For a given Nb value, draw n separate times from a multinomial distribution with the probability vector given by the barcode lineage frequencies in the donor sample and the number of trials being Nb. A given draw can be considered a random realization of barcode lineages in a recipient sample seeded by a bottleneck size of Nb. We let n = 500; higher values of n did not alter the results.
  b. For each of the n draws from the multinomial distribution, quantify the number of barcode lineages present in the recipient sample.
  c. From the n independent draws, calculate the mean and standard deviation of the number of barcode lineages present in the recipient sample.
  d. Calculate the probability of observing the observed number of barcodes in the recipient using a normal distribution with mean and standard deviations as calculated above. The log of this probability yields the log-likelihood of the bottleneck size being Nb.
4. Identify the maximum likelihood estimate of Nb as the Nb yielding the highest log-likelihood. Calculate the 95% confidence interval of Nb as the set of Nb values that yield log-likelihood values within 1.92 log-likelihood units of the maximum likelihood estimate of Nb.

## ACKNOWLEDGEMENTS

We thank Jesse Bloom and Yoshihiro Kawaoka for providing key plasmids and reagents. We thank Christopher Brooke and Brigette Martin for assistance in virus rescue protocols. This work is funded by AI125392 to AM and TCF and by the National Institute of Allergy and Infectious Diseases under HHS contract HHSN27220140006C for the St. Jude Center of Excellence for Influenza Research and Surveillance, NIH grant R01AI140766, and ALSAC to SSC. KAA is supported by GM007215. KMB is supported by AI145182. LAH is supported by HG002760. GAS is supported by a Rath Foundation Wisconsin Distinguished Graduate Fellowship and AI007414. AM holds an Investigators in the Pathogenesis of Infectious Disease Award from the Burroughs Wellcome Fund.

