## Supplemental Figures 1-7 for "Influenza A virus undergoes compartmentalized replication *in vivo* dominated by stochastic bottlenecks"

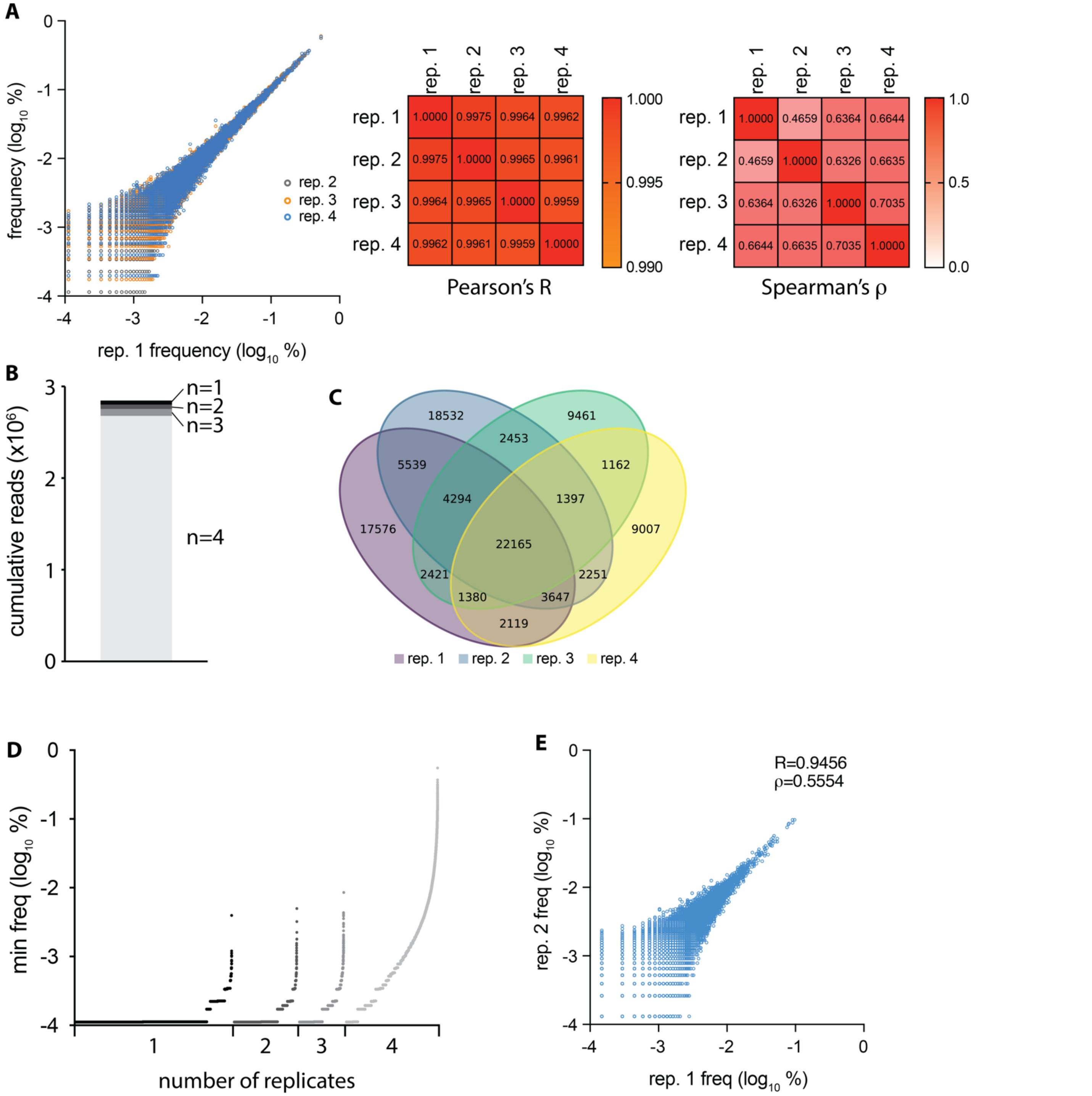


**Supplemental Figure 1. Replicate sequencing of viral stock with an NheI registration mark highlights reproducibility of barcode enumeration. Associated with Figure 3.** A) The barcodes present on HA in virus stock with the NheI registration mark were subject to 4 replicate sequencing runs to assess reproducibility. The frequency of individual barcodes in replicates 2-4 are plotted relative to their frequency in replicate 1 (left). Pearson's R and Spearman's ρ correlation coefficients calculated between all replicate pairs (right). B) Almost all HA barcode sequencing reads are shared in all for replicates. Sequence reads were grouped based on their presence in n = 1, 2, 3 or 4 replicates. C) Venn diagram highlighting overlap of HA barcode identity in replicate sequencing, independent of the abundance of any individual barcode. D) Replicate sequencing of HA barcodes reproducibly enumerates all but the lowest frequency barcodes. Barcodes were separated based on their presence in 1, 2, 3 or 4 replicates. The minimum frequency of an individual barcode across all replicates was plotted. Barcodes with the lowest frequency tend to appear in only a subset of replicates. E) Replicate sequencing of PA barcodes present in virus stock with the NheI registration mark. R = Pearson's correlation coefficient. ρ = Spearman's rank correlation coefficient.

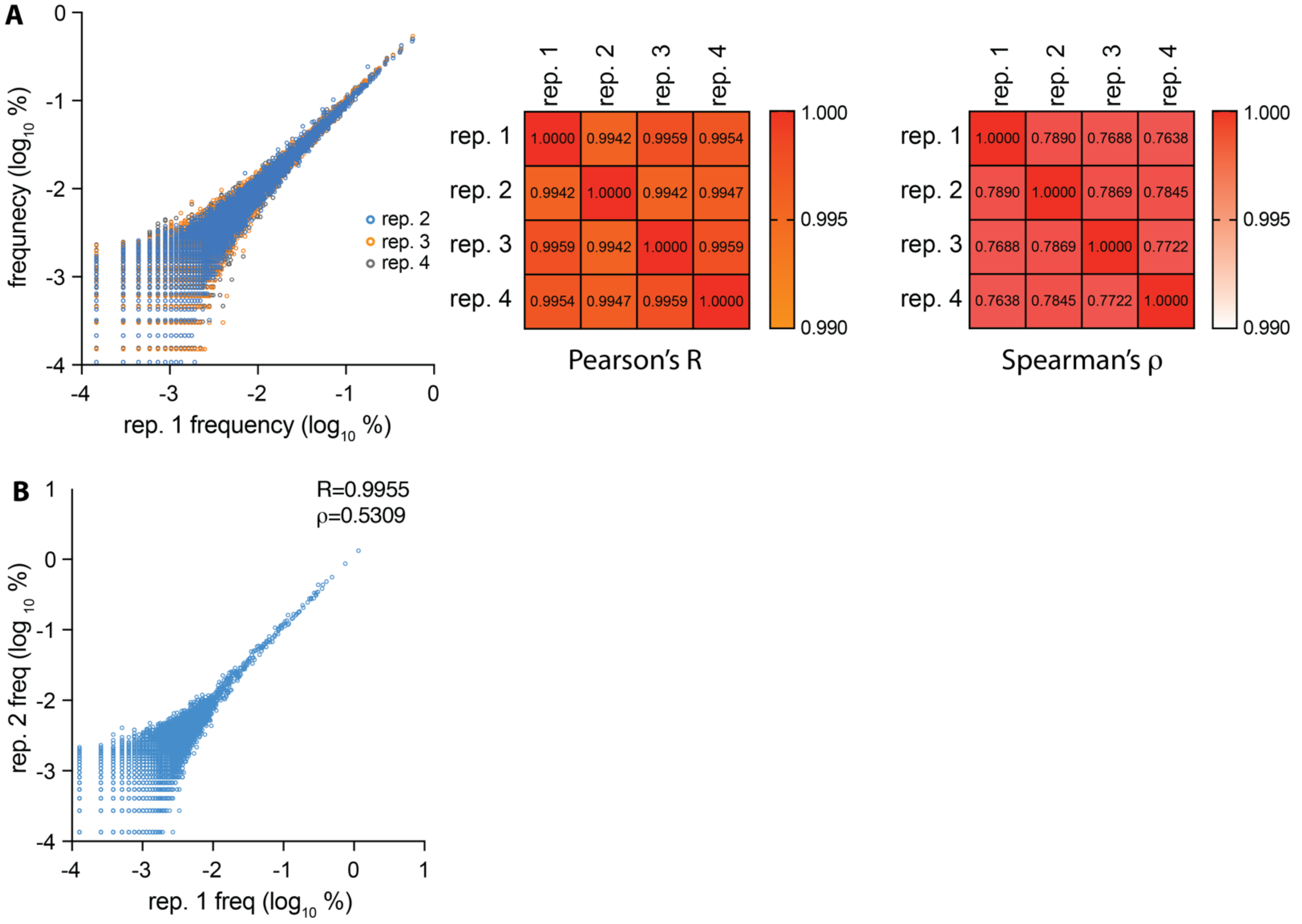


**Supplemental Figure 2. Replicate sequencing of viral stock with a PstI registration mark highlights reproducibility of barcode enumeration. Associated with Figure 3.** A) The barcodes present on HA in virus stock with the PstI registration mark were subject to 4 replicate sequencing runs to assess reproducibility. The frequency of individual barcodes in replicates 2-4 are plotted relative to their frequency in replicate 1 (left). Pearson's R and Spearman's ρ correlation coefficients calculated between all replicate pairs (right). B) Replicate sequencing of PA barcodes present in virus stock with the PstI registration mark. R = Pearson's correlation coefficient. ρ = Spearman's rank correlation coefficient.

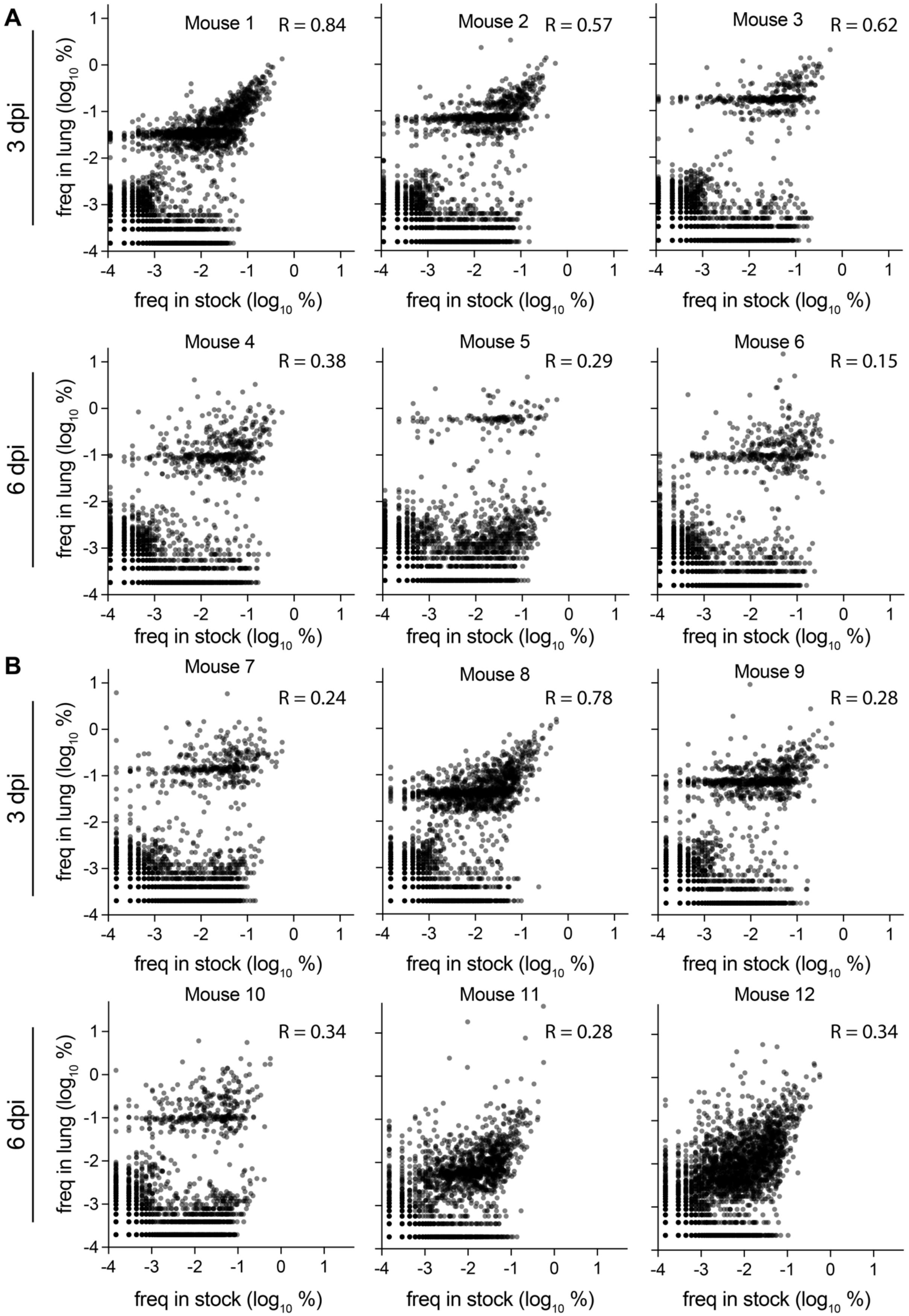


**Supplemental Figure 3. Diverse populations in mouse lungs are partially correlated with high frequency members in the inoculum at early times during infection. Associated with Figure 4.** A) The frequency of lineages in the inoculum was compared to those in mice inoculated with HA-K153E-NheI libraries at 3 and 6 dpi. B) Same as A for the HA-K153E-PstI libraries. R = Pearson's correlation coefficient.

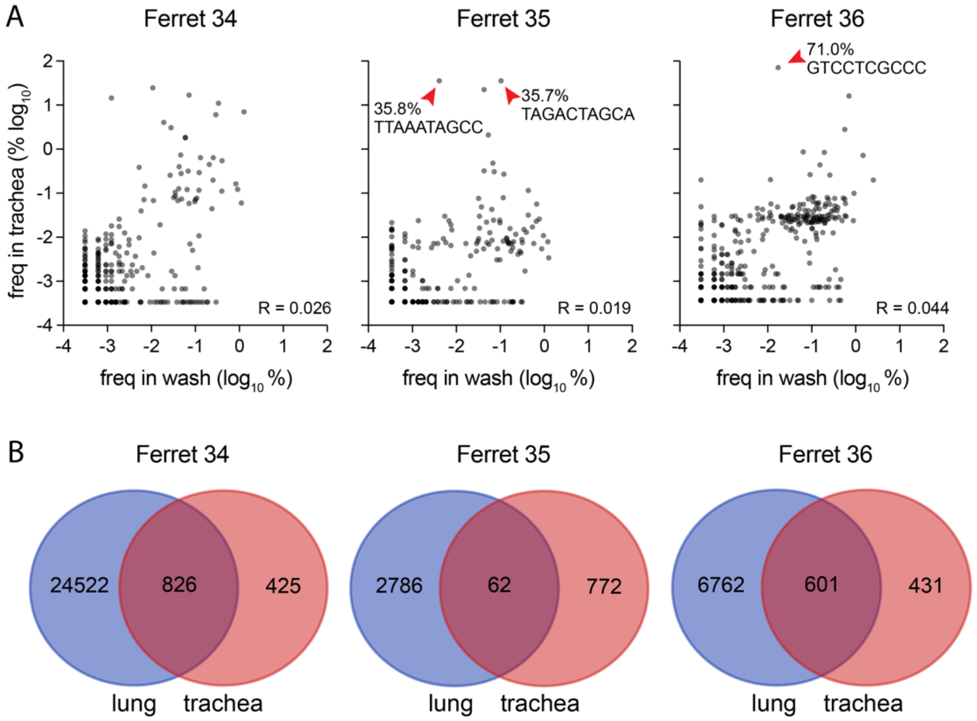


**Supplemental Figure 4. Replication of reduced populations in the trachea. Associated with Figure 5.** A) HA barcode frequencies were compared between the nasal wash (3 dpi) and trachea (5 dpi) for ferret 34, 35, and 36. Red arrowheads highlight dominant barcodes with frequency > 30%. R = Pearson's correlation coefficient. Note that barcodes unique to the nasal wash or trachea are not plotted here. B) Venn diagram of lineages present in the trachea and all lung lobes at 5 dpi.

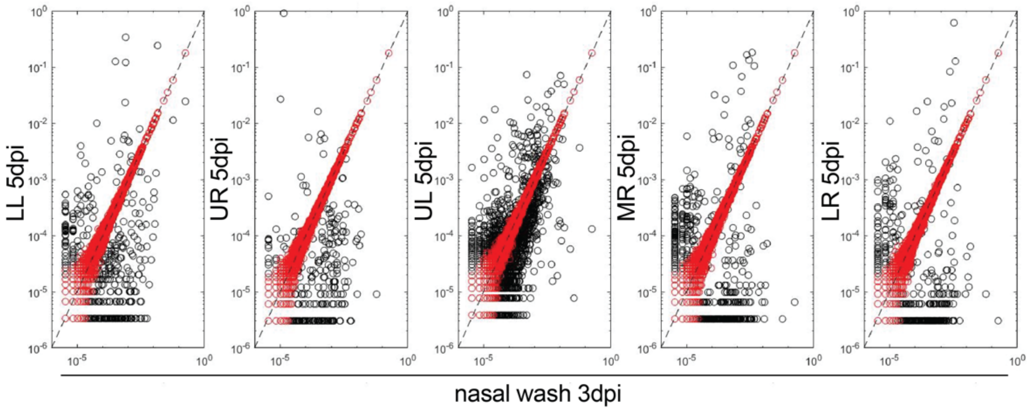


**Supplemental Figure 5. Modelling suggests tighter bottlenecks in animals. Associated with Figure 6.** Maximum likelihood estimates of bottleneck size was used to forward simulate a mock dataset where the starting population in nasal washes from ferret 34 at 3 dpi passes through the bottleneck yielding predicted frequencies in the lung lobe. Modeled data (red) show a strong correlation between frequency in the donor and recipient populations, whereas actual data from ferret 34 (black) show large differences in frequencies indicative of tighter bottlenecks.

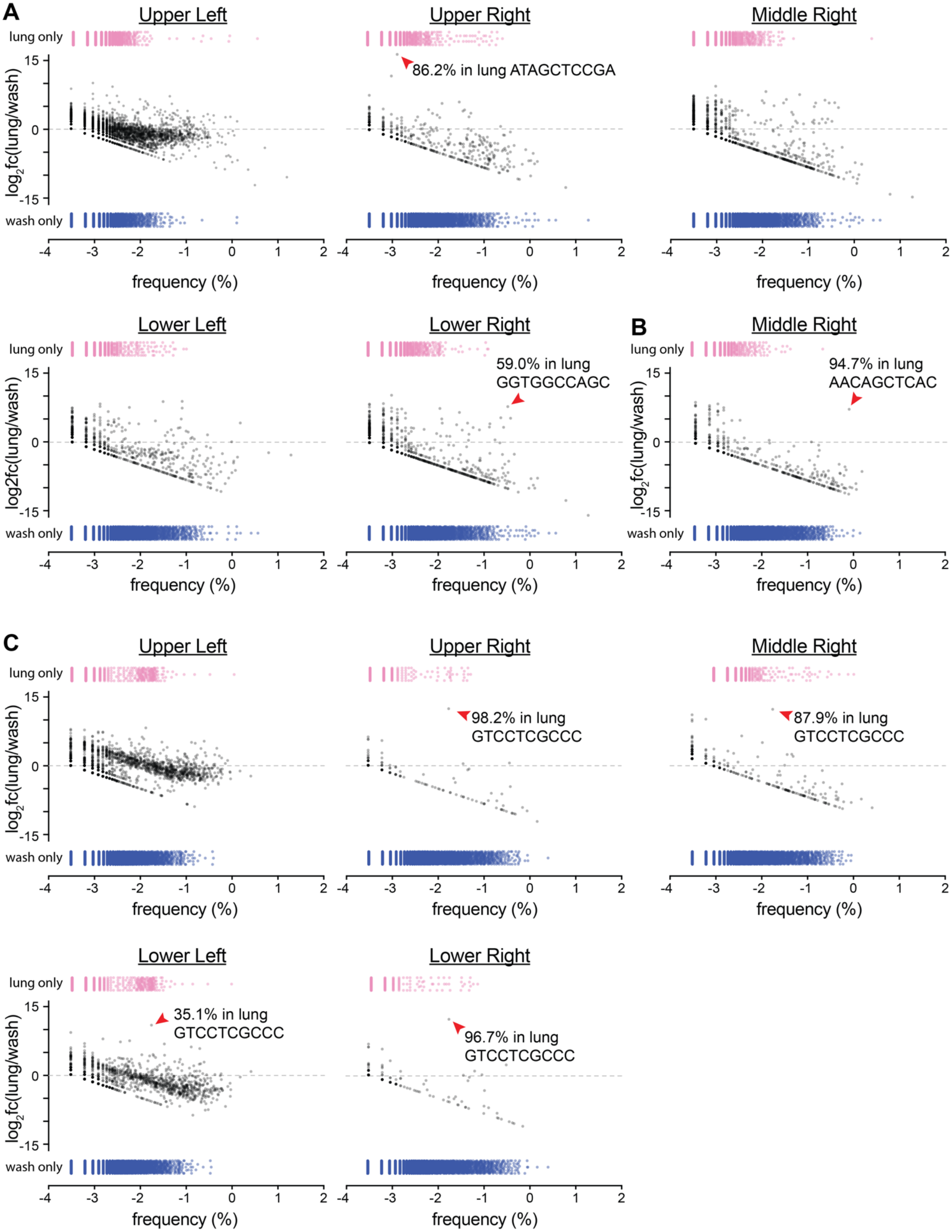


**Supplemental Figure 6. Bottlenecks result in stochastic outgrowth of viruses in lung lobes. Associated with Figures 6-7.** HA barcode frequencies were determined in the nasal wash (3 dpi) and lung lobes (5 dpi) for A) ferret 34, B) ferret 35, and C) ferret 36. The $\log_2$-fold change ($\log_2$fc) in frequency between the lung and the nasal wash was plotted as a function of barcode frequency in the nasal wash (black dots). Barcodes present only in the nasal wash (blue) or lung lobe (pink) are plotted as a function of their frequency within their respective populations. Red arrowheads highlight dominant barcodes with frequency > 30%.

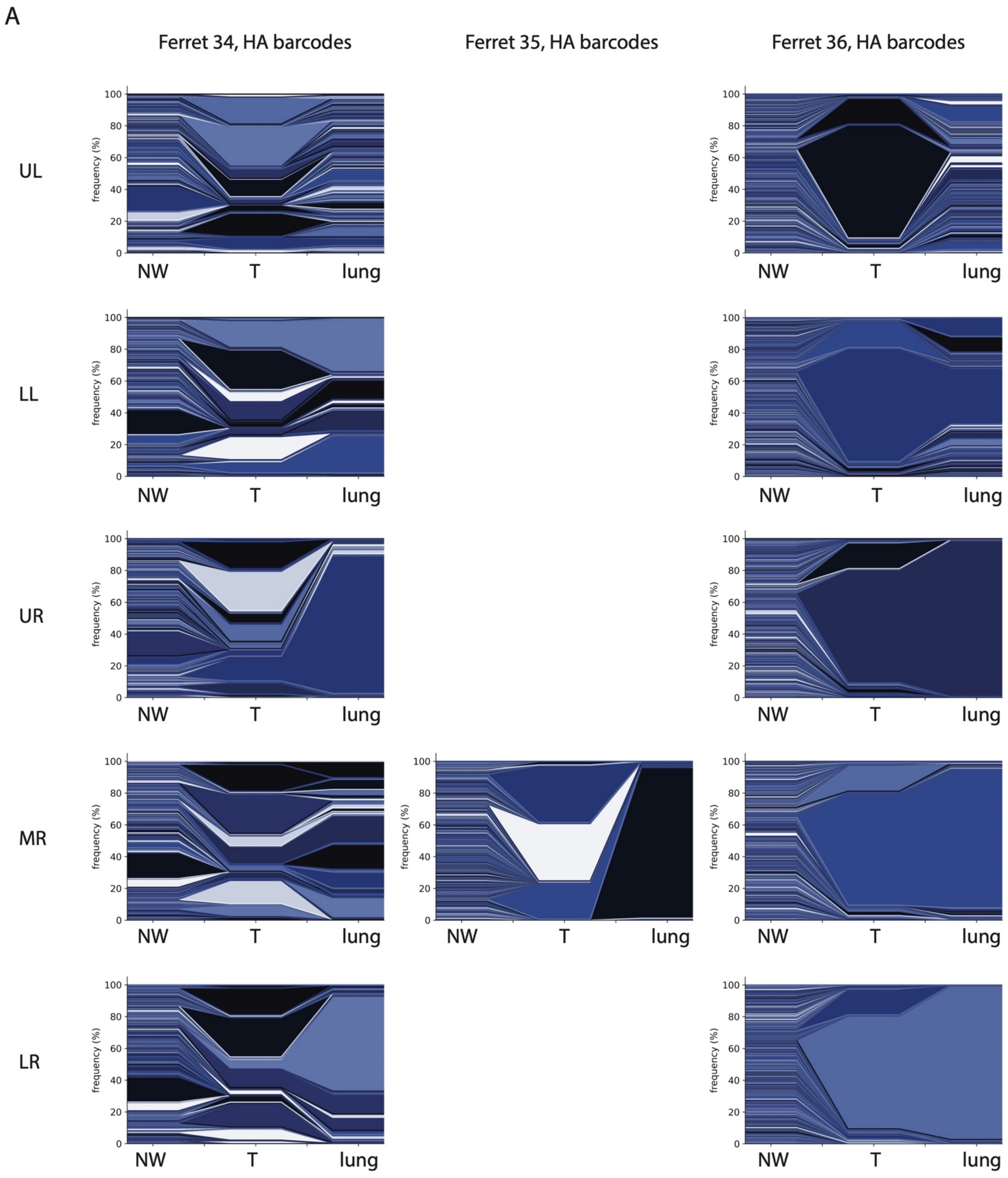
A
Ferret 34, HA barcodes
Ferret 35, HA barcodes
Ferret 36, HA barcodes
UL
LL
UR
MR
LR
frequency (%)
NW
T
lung

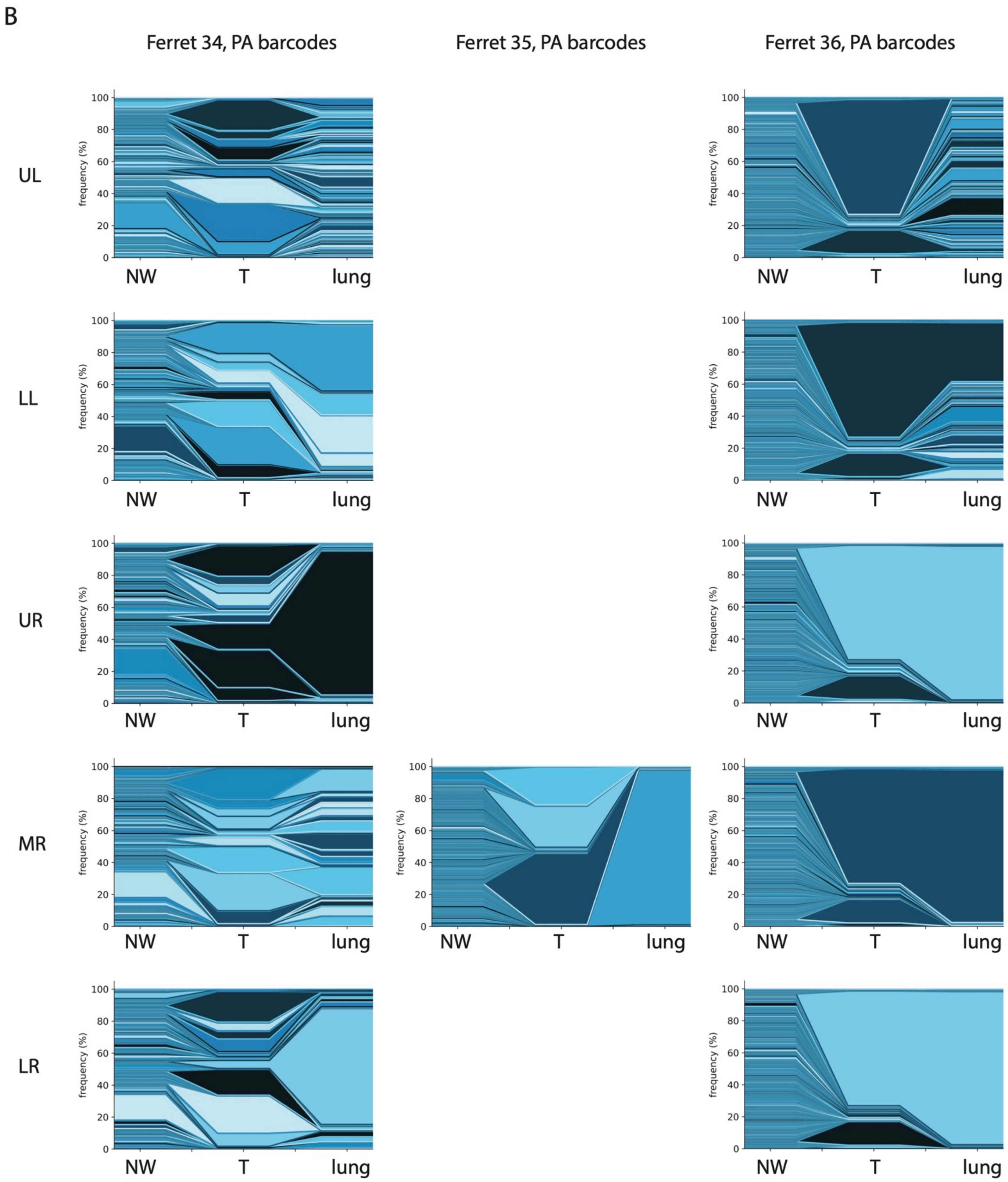


**Supplemental Figure 7. Tracking barcodes throughout the respiratory tract. Associated with Figures 5-7.** Plots are shown for all lung lobes in which infectious virus was detected. Barcode frequency was plotted to illustrate migration from the upper respiratory tract into distinct lung lobes for (A) HA and (B) PA. Each color represents a unique barcode. Colors are not conserved across samples. NW = nasal wash, 3 dpi. T = trachea. Lung = lung lobe indicated by row.
